# SILAC proteomics implicates the ubiquitin conjugating enzyme UBE2D in SOCS1-mediated downmodulation of the MET receptor in hepatocytes

**DOI:** 10.1101/2020.07.31.231225

**Authors:** Madanraj Appiya Santharam, Akhil Shukla, Awais Ullan Ihsan, Maryse Cloutier, Dominique Levesque, Sheela Ramanathan, François-Michel Boisvert, Subburaj Ilangumaran

## Abstract

Suppressor of Cytokine Signaling 1 (SOCS1) functions as a tumor suppressor in hepatocellular carcinoma (HCC) and many other types of cancers. SOCS1 mediates its functions by inhibiting tyrosine kinases, promoting ubiquitination and proteasomal degradation of signal transducing proteins, and by modulating transcription factors. Here, we studied the impact of SOCS1 on the hepatocyte proteome using Stable Isotopic Labelling of Amino acids in Cell culture (SILAC)-based mass spectrometry on the Hepa1-6 murine HCC cell line stably expressing wildtype SOCS1 or a mutant SOCS1 with impaired SH2 domain. As SOCS1 regulates the hepatocyte growth factor (HGF) receptor MET, the SILAC-labelled cells were stimulated or not with HGF. Following mass spectrometry analysis, differentially modulated proteins were identified, quantified and analyzed for pathway enrichment. Of the 3440 proteins identified in Hepa-SOCS1 cells at steady state, 181 proteins were significantly modulated compared to control cells. The SH2 domain mutation and HGF increased the number of differentially modulated proteins. Protein interaction network analysis revealed enrichment of SOCS1-modulated proteins within multiprotein complexes such as ubiquitin conjugating enzymes, proteasome, mRNA spliceosome, mRNA exosome and mitochondrial ribosome. These findings suggest that SOCS1, induced by cytokines, growth factors and diverse other stimuli, may dynamically modulate of large macromolecular regulatory complexes to help maintain cellular homeostasis. Notably, the expression of UBE2D ubiquitin conjugating enzyme, which is implicated in the control of growth factor receptor tyrosine kinase signaling, was found to be regulated by SOCS1.

## Introduction

Suppressor of cytokine signaling 1 (SOCS1) is a negative feedback regulator of the JAK-STAT signaling pathway induced by cytokines and is essential to control interferon gamma signaling [1]. SOCS1 also regulates receptor tyrosine kinases (RTK) activated by growth factors such as the hepatocyte growth factor (HGF) [2]. SOCS1 deficiency accelerates hepatocyte proliferation during liver regeneration [3]. Loss of SOCS1 gene expression in the liver has been reported to occur in a high proportion of hepatocellular carcinoma (HCC) specimens [4,5], and SOCS1-deficient mice display increased susceptibility to experimental hepatocarcinogenesis [4–7]. These findings and epigenetic repression of the *SOCS1* gene in many other cancers support a tumor suppressor role of SOCS1 [8]. Understanding the molecular mechanisms of tumor suppression mediated by SOCS1 is an active area of research as it could lead to the identification of druggable molecular targets.

Studies on SOCS1-deficient mice and hepatocyte cell line models have shown that SOCS1 can control the development and progression of HCC via at least by two mechanisms: (i) attenuation of hepatocyte growth factor (HGF)-induced MET receptor tyrosine kinase (RTK) signaling [3,9], and (ii) inhibition of the aberrant oncogenic potential of the cyclin dependent kinase inhibitor CDKN1A [7]. The oncogenic potential of CDKN1A results from its cytosolic retention caused by AKT-dependent phosphorylation downstream of deregulated growth factor signaling [7,10]. It is possible that SOCS1 may also regulate other growth promoting and potentially oncogenic signaling pathways that are implicated in HCC pathogenesis [11,12]. Even though SOCS1 was discovered as a negative feedback regulator of IL-6-induced JAK-STAT signaling [13], this pathway did not show deregulation in SOCS1-deficient livers [3,7], possibly because SOCS3 is necessary and sufficient to attenuate IL-6-induced STAT3 activation in hepatocytes [14,15]. Similar complementation by SOCS3 may mask at least some of the oncogenic signaling pathways activated by the loss of SOCS1.

SOCS1 has been shown to interact with diverse signaling molecules in different cell types [16,17], lending support to the idea that SOCS1 may regulate other growth promoting signaling pathways in hepatocytes. SOCS1 contains a central SH2 domain, which binds phosphorylated tyrosine residues in activated signaling molecules such as JAK and MET kinases [3,18,19]. Poly-proline residues in the amino terminal region of SOCS1 can interact with a number of SH3 domain containing proteins such as GRB2, p85 regulatory subunit of PI3K (PIK3R1) and NCK [20]. The SOCS box in the carboxy terminus of SOCS1, which is composed of a BC box and a cullin box, connects with elongin B, elongin C and Cullin5 that interact with the RING-finger protein RBX2 to assemble the protein ubiquitin ligase CRL^SOCS1^ [21,22]. Many of the SOCS1-interacting proteins, including JAKs, MET and CDKN1A undergo SOCS1-dependent ubiquitination and proteasomal degradation [7,9,23–25]. Recent mass spectrometry studies on SOCS1-interacting proteins in myeloid leukemia cells identified many previously known and new protein targets [17].

While the above functions of SOCS1 occur in the cytosol, several reports indicate a regulatory role for SOCS1 in the nucleus as well. SOCS1 possesses a nuclear localization signal (NLS) and regulates NF-kB signaling by promoting ubiquitination and degradation of p65RelA within the nucleus [26–29]. SOCS1 lacking the NLS was shown to be defective in suppressing innate interferon-induced activation of gene promoters [30]. We have shown that SOCS1 modulates the transcriptional activation of p53 by promoting its phosphorylation by ATM-ATR kinases [31]. Recent studies indicate that the tumor suppressor activity mediated via SOCS1-p53 axis is inhibited by tyrosine phosphorylation of SOCS1 by oncogenic SRC kinase signaling [32]. Indeed the p53 motif that interacts with Tyr-phosphorylated SOCS1 is also present in several other transcription factors such as STAT, CEBPZ and FOXM1, and hence they could also be potentially be regulated by SOCS1. Based on these premises, we hypothesized that even though SOCS1 is only transiently induced by cytokines, growth factors and diverse other stimuli, it can potentially modulate the cellular protein landscape either directly via protein ubiquitination or indirectly via modulating transcription factors.

In order to systematically characterize the molecular targets of SOCS1-dependent regulation of hepatocarcinogenesis, we carried out an unbiased differential proteomic approach on a murine HCC cell line Hepa1-6. It has been shown that stable expression of SOCS1 in Hepa1-6 cells markedly reduces its growth as subcutaneous tumor in syngeneic C57BL6 mice, and attenuates HGF-induced cell proliferation, migration and invasion [9,33]. The present study was undertaken to characterize proteins regulated by SOCS1 in Hepa cells using the ‘stable isotope labeling with amino acids in cell culture’ (SILAC)-based quantitative mass spectrometry [34]. Our findings show that constitutive expression of SOCS1 not only downmodulates several proteins but also upregulates a number of proteins at steady state, and analysis of their relationship revealed prominent clustering within multiprotein regulatory complexes such as proteasome and exosome involved in the processing of proteins and RNA, respectively. Our findings suggest that SOCS1 may be involved in modulating macromolecular complexes that regulate cellular homeostasis in hepatocytes.

## Materials and Methods

### Cell lines

Hepa1-6 cells obtained from ATCC (CRL-1830) were transduced with pWPT lentiviral vector containing wildtype SOCS1 (Hepa-SOCS1, HS) or SOCS1 with R105K mutation within the SH2 domain (Hepa-SOCS1R105K, HR) or the empty vector (Hepa-vector, HV). All three cell lines were grown in DMEM containing 5% Fetal Bovine Serum (FBS).

### Stable isotope labeling with amino acids in cell culture (SILAC)

Cells were resuspended in Arg and Lys-free DMEM (Wisent, #319-118-CL) supplemented with 5% dialyzed FBS (Gibco, catalogue #26400-044), 2mM GlutaMax (Gibco, #35050061) and 100 U/ml Penicillin/streptomycin (Wisent, #450-201-EL). To this medium, the following combinations of L-Arg and L-Lys carrying different carbon (C), nitrogen (N) and deuterium (D) isotopes were added: (i) light (L: R_0_K_0_) isotopes (^12^C^14^N on both Arg and Lys; Sigma-Aldrich: A6969, L8662), (ii) medium (M: R_6_K_4_ = +10Da) isotopes (^13^C on Arg, 4 D on Lys; Cambridge isotope laboratories, L-Arg: catalogue #CLM-2265-H-PK; L-Lys: #DLM-2640-PK) or (iii) heavy (H: R_10_K_8_ = +18Da) isotopes (^13^C^15^N on both Arg and Lys: Cambridge isotope laboratories, L-Arg: #CNLM-539-H-PK; L-Lys: #CNLM-291-H-PK). HV, HS and HR cells were seeded in 100 mm petri dishes (25 x10^3^ cells/ dish) in SILAC media with light, medium or heavy isotope-labelled amino acids, and cultured for 6-8 days to allow incorporation of the labelled amino acids into proteins over more than 5 population doublings. Medium was replenished after 3 days and then every 2 days. The cells were switched to medium containing 0.5% dialyzed FBS and the respective isotope-labelled amino acids. After overnight serum starvation, one set of the three cultures were stimulated with hepatocyte growth factor (HGF) (Peprotech, #315-23; 25ng/ml) for 24h, and the other set was left unstimulated and lysed at the same time as HGF-stimulated cells.

### Cell lysis, protein separation and mass spectrometry

The peptides were prepared for mass spectrometry as previously described [35]. The cells were thoroughly washed in PBS before lysis in radioimmunoprecipitation assay (RIPA) buffer (50mM Tris-HCl, pH 7.4, 1% Triton-X-100, 0.5% sodium deoxycholate, 0.1% SDS, 150mM NaCl, 2mM EDTA, 50mM sodium fluoride; 250μl/petri dish). The cells were collected using a cell scrapper, incubated on ice for 30 min and sonicated using a probe sonicator (Branson 150D Ultrasonic Cell Disruptor, Danbury, CT). The lysates were centrifuged at 12,000g for 20 mins to remove cell debris and protein concentration in supernatants was determined using DC™ Protein Assay (Bio-Rad, #5000111). Twenty μg of proteins from L, M and H isotope-labeled cultures from three experiments were pooled as shown in the Figure 1A. The protein mixtures were separated by SDS-PAGE electrophoresis, and each lane was cut into 4 pieces. The gel stain was removed by repeated washing in acetonitrile (CH_3_CN; Fluka, #34967) and 20mM ammonium bicarbonate (NH4HCO3; Fluka, #09830), followed by overnight *in situ* trypsin digestion (Pierce, #90058; 12.5ng/ml trypsin in 50mM acetic acid, 20mM sodium bicarbonate). The peptides were extracted into the trypsin digestion by using equal amount of CH_3_CN and incubating for 30 min at 30°C. Following collection of the supernatant, the same step was repeated with 1% formic acid. These two steps were repeated until the gel pieces were completely dehydrated. The solutions containing the peptides were dried using a Speedvac concentrator (Eppendorf™ Vacufuge™ Concentrator, #07-748-13) and suspended in 0.1% TFA (Sigma-Aldrich, #T6508). The peptides were cleaned from other impurities using ZipTips (Millipore, # ZTC18S) and re-suspended in 1% formic acid (Fisher, #A117-50). These purified peptides were quantified using a nanodrop (Thermo Fisher Nanodrop 2000c) before mass spectrometry.

**Fig. 1.**
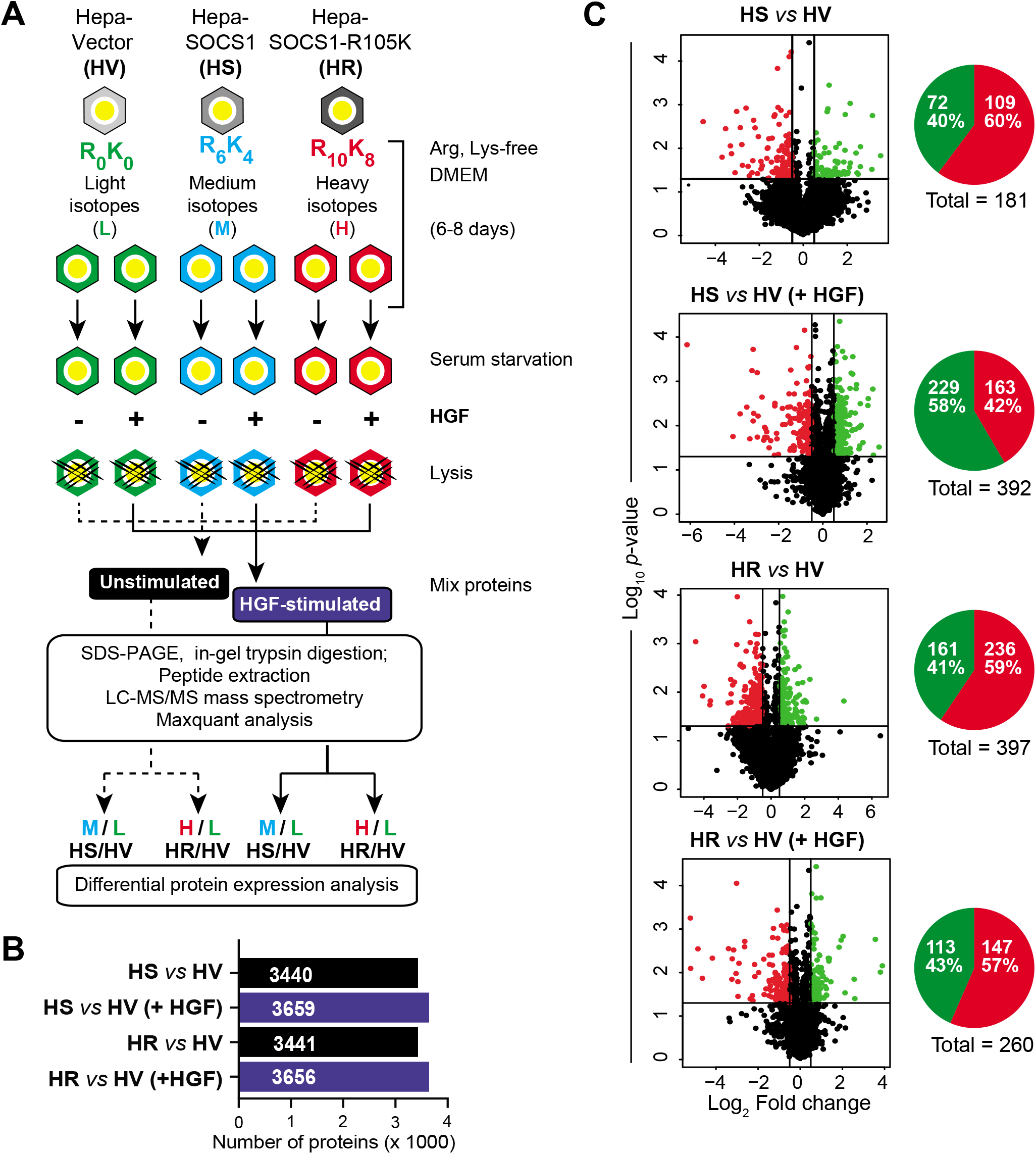
Identification of proteins differentially regulated by SOCS1 in hepatocytes. (A) The study protocol. HV, HS and HR cells were grown in SILAC medium containing 5% dialyzed FBS and different isotopes of Arg and Lys, namely light (R_0_K_0_), medium (R_6_K_4_) and heavy (R_10_K_8_), for 5 generations. At ~60-70% confluent level, the cultures were serum starved (0.5% dialysed serum) and stimulated with mouse HGF (25 ng/ml) for 24 h (+ HGF) or left unstimulated at steady state. Cells were lysed in SDS-PAGE sample buffer and 20μg of proteins for each condition from three experiments were mixed into two combinations, representing unstimulated and HGF-stimulated cells, as shown. Proteins were separated on SDS-PAGE gels that were sliced and peptides extracted following in-gel trypsin digestion. Eluted peptides were analysed by LC-MS/MS and proteins identified from the mass data in RAW files using the MaxQuant software. (B) Peptides that differed in their molecular mass by 10 Da (HS versus HV) or 18 Da (HR versus HV), which reflects the incorporated isotope-labelled Arg and Lys, identified the total number of proteins in each indicated comparison. (C) Proteins that are significantly modulated by SOCS1 or SOCS1R105K were identified by plotting the fold change in protein expression levels from triplicate experiments for each indicated comparison against the log_10_ of *p*-value, which was calculated for each protein by unpaired *t*-test using an R software package. Volcano plots (left column) are used to identify differentially expressed proteins (*p*<0.05 and fold change 0.5, i.e., 200% or 50%). Upregulated and downregulated proteins, identified as green and red color dots in each plot, respectively. Pie charts (right) show the total number of significantly upregulated (green) and downnregulated (red) proteins in each comparison.

### LC-MS/MS

A total of 2 μg of trypsin-digested peptides were separated using a Dionex Ultimate 3000 nanoHPLC system. The sample was loaded with a constant flow of 4 μl/min onto a Trap column (Acclaim PepMap100 C18 column ((0.3 mm id x 5 mm, Dionex Corporation)). The peptides were then eluted onto an analytical column (PepMap C18 nano column (75 μm x 50 cm, Dionex Corporation)) with a linear gradient of 5-35% solvent B (composed of 90% acetonitrile with 0.1% formic acid) over four hours with a constant flow of 200 nl/min. The HPLC was coupled to an OrbiTrap QExactive mass spectrometer (Thermo Fisher Scientific Inc.) via an EasySpray source, with a spray voltage set to 2.0 kV and the column set at 40°C. Full scan MS survey spectra (m/z 350-1600) were acquired in profile mode at a resolution of 70,000 after accumulation of 1,000,000 ions. The ten most intense peptide ions from the preview scan were selected for fragmentation by collision induced dissociation (CID, with a normalized energy set at 35% and the resolution set at 17,500) after the accumulation of 50,000 ions, with a filling time of 250 ms for the full scans and 60 ms for the MS/MS scans. Screening of the precursor ion charge state was enabled and all unassigned charge states as well as singly, 7 and 8 charged species were rejected. A dynamic exclusion list was allowed to a maximum of 500 entries with a retention time of 60 seconds, using a relative mass window of 10 ppm. The lock mass option was enabled for survey scan to improve mass accuracy. Data were acquired using the Thermo Scientific™ Xcalibur™ software. The mass spectrometry proteomics data have been deposited to the ProteomeXchange Consortium via the PRIDE [36] partner repository with the dataset identifier PXD019135.

### Identification and Analysis of proteins

The peptides were identified and quantified using MaxQuant software version 1.5.2.8 [37] with the reference database from Uniprot KB (*Mus musculus*). The proteins obtained were filtered by removing the known possible contaminants, reverse sequences and proteins identified with a single peptide from the list. The fold change for each protein was obtained by using the formula log to the base 2 of protein ratios of each respective combination. The significantly modulated proteins were screened using an R analysis software (R studio) by performing a one-sample T-test for each protein using the fold change. A criterion of 5% false discovery rate (FDR) was applied along with the restriction of more than 2 peptides identified for each protein to be considered as a confident hit. The list was further shortened by using a minimum cut-off of 0.5-fold change (FC) compared to control.

### Pathway analysis and interaction profiling

Proteins that are differentially expressed in each set of comparisons was loaded in the PANTHER database (http://www.pantherdb.org) to carry out enrichment analysis of protein classes and signaling pathways. A minimum representation of three (for HS *vs* HV) or four proteins (for all other comparisons) was considered for signaling pathway enrichment. Shared proteins in different comparisons were identified using the Venny 2.1 (https://bioinfogp.cnb.csic.es/tools/venny/) and Biovenn (http://www.biovenn.nl) [38] softwares. Protein interaction profiling was done using the STRING database (https://string-db.org) [39] and illustrations were made using Cytoscape software version 3.7.1 [40].

### Correlation between SOCS1 and select genes in HCC

The gene expression analysis was performed on the RNAseq data from the TCGA provisional dataset on liver HCC (LIHC) generated by the TCGA Research Network (https://www.cancer.gov/tcga) [41]. The provisional TCGA-LIHC cohort contains 442 specimens, of which RNAseq V2 data on *SOCS1, UBE2D* variants, *MET* and *CBL* are available for 348 samples. The gene expression dataset was accessed via the cBioportal suite for cancer genomics research (https://www.cbioportal.org) to evaluate co-expression.

### cDNA preparation and qRT-PCR

After washing the cells with PBS, total RNA from Hepa-V, Hepa-SOCS1 and Hepa-SOCS1R105K was extracted using RiboZol™ (AMRESCO, Solon, OH). After quality control by UV absorption, complementary DNA strand was synthesized from one μg of total RNA using QuantiTect^®^ reverse transcription kit (Qiagen). Quantitative RT-PCR gene amplification was carried out using the CFX-96 thermocycler (Bio-Rad, Mississauga, ON) using the primers listed in Supplementary Table S1A. Fold induction of specific genes was calculated by comparing the expression levels of housekeeping gene as a reference.

### Western Blot

Hepa-S, Hepa-R and Hepa-V cells were grown to 70% confluency in standard 60mm petri dishes, serum starved for 12-16h and exposed to mouse HGF (25ng/ml) with or without cycloheximide (CHX, 100μM) for different time periods. Cells were washed thoroughly in PBS and lysed in RIPA lysis buffer, collected in microcentrifuge tubes, vortexed for 30 seconds, left on ice for 15 min and centrifuged at 15000g for 20 min to remove cell debris. Protein concentration was determined using Pierce™ BCA Protein Assay Kit (Thermo Fisher, #23227), 10-15 μg proteins were separated on SDS-PAGE gels and western blots were probed with the primary antibodies listed in Supplementary Table S1B. Secondary antibodies as well as ECL (enhanced chemiluminescence) reagents were obtained from GE Healthcare Life Sciences (Pittsburg, PA, United States). Western blot images were captured on the ChemiDoc™ MP System (Bio-Rad) and densitometry quantification was carried out using Adobe Photoshop CC 2020 software.

## Results

### SOCS1 modulates the expression of hundreds of hepatocyte proteins

To carry out a systematic analysis of proteins modulated by SOCS1 in hepatocytes, we used Hepa1-6 murine HCC cell line expressing SOCS1 (Hepa-SOCS1; HS) or its SH2 domain mutant SOCS1R10K (Hepa-SOCS1R105K; HR) from a stably integrated lentiviral vector [3]. The functionality of these two exogenous proteins was verified by exposing HS, HR and control cells expressing the vector backbone (Hepa-vector; HV) briefly to IFNγ and evaluating STAT1 phosphorylation at different time points after IFNγ withdrawal (Supplementary Fig. S1A). HV cells showed marked phosphorylation of STAT1 that gradually diminished over time. The IFNγ-induced STAT1 phosphorylation was almost completely abrogated in HS cells, whereas STAT1 phosphorylation in HR cells was comparable to HV cells, albeit slightly delayed and prolonged (Supplementary Fig. S1A). Stable expression of SOCS1 significantly reduced cell growth that was evident only after 6 days of culture (Supplementary Fig. S1B). The expansion of HR cells was comparable to that of HV cells until 7 days.

HV, HS and HR cells grown in DMEM-5% FBS were transferred to SILAC medium (DMEM containing 5% dialyzed serum) containing different mass isotopes as indicated in Fig. 1A. After 5-6 days of culture to maximize the incorporation of the mass label, the cells were serum starved overnight and triplicate cultures were stimulated with HGF 24h. Lysates of control and HGF-stimulated cells were mixed in the order shown in Fig. 1A, proteins separated on SDS-PAGE and digested *in situ* in gel slices, peptides analyzed by LC-MS/MS, and differentially expressed proteins were quantified and identified by MaxQuant for the indicated pairwise comparisons. All the four different comparisons namely, HS versus HV at steady state (HS *vs* HV), HS versus HV stimulated with HGF (HS *vs* HV (HGF)), HR versus HV at steady state (HR *vs* HV) and HR versus HV stimulated with HGF (HR *vs* HV (HGF)) identified similar numbers (~3,500) of proteins (Fig. 1B). The fold change was plotted against the *p*-value of differential expression in Volcano plots to select proteins that were reduced by 50% or increased by 2-fold and with significant *p* values of <0.05. This analysis identified 109 downregulated and 72 upregulated proteins in HS cells compared to HV cells (Fig. 1C). These numbers almost doubled following HGF stimulation but with more upregulated (229) proteins than downregulated (163) proteins. Surprisingly, HR cells showed more downregulated and upregulated proteins than HS cells but without the inversion in their numbers following HGF stimulation as seen in HS cells. All downregulated and upregulated proteins within each group are listed in Supplementary Tables S2A-S2D along with fold change and *p*-values.

### SOCS1 modulates diverse classes of cellular proteins in various signaling pathways

To characterize the proteins modulated by SOCS1, the significantly downregulated and upregulated proteins (FDR *p*<0.05) within each group were combined and analyzed using the PANTHER database. Analysis of the protein families modulated by SOCS1 revealed diverse classes of proteins (Fig. 2A) that showed considerable level of overlap among the four comparison groups. Among them, nucleic acid binding proteins occupied the top rank in cells expressing either SOCS1 or SOCS1R105K at steady state, but this class descended in order in HGF-stimulated cells. Transcriptions factors also figured predominantly in all four groups. Transporters and proteins involved in membrane traffic are the two other groups that were modulated by SOCS1 proteins in all groups. Strikingly, signaling molecules and ligases were far down the list of proteins modulated by SOCS1. Molecules involved in defense and immunity were found at the lower end of these lists. Conversely, hydrolases and enzyme modulators, which include a large number of proteins, showed an inverse trend.

**Fig. 2.**
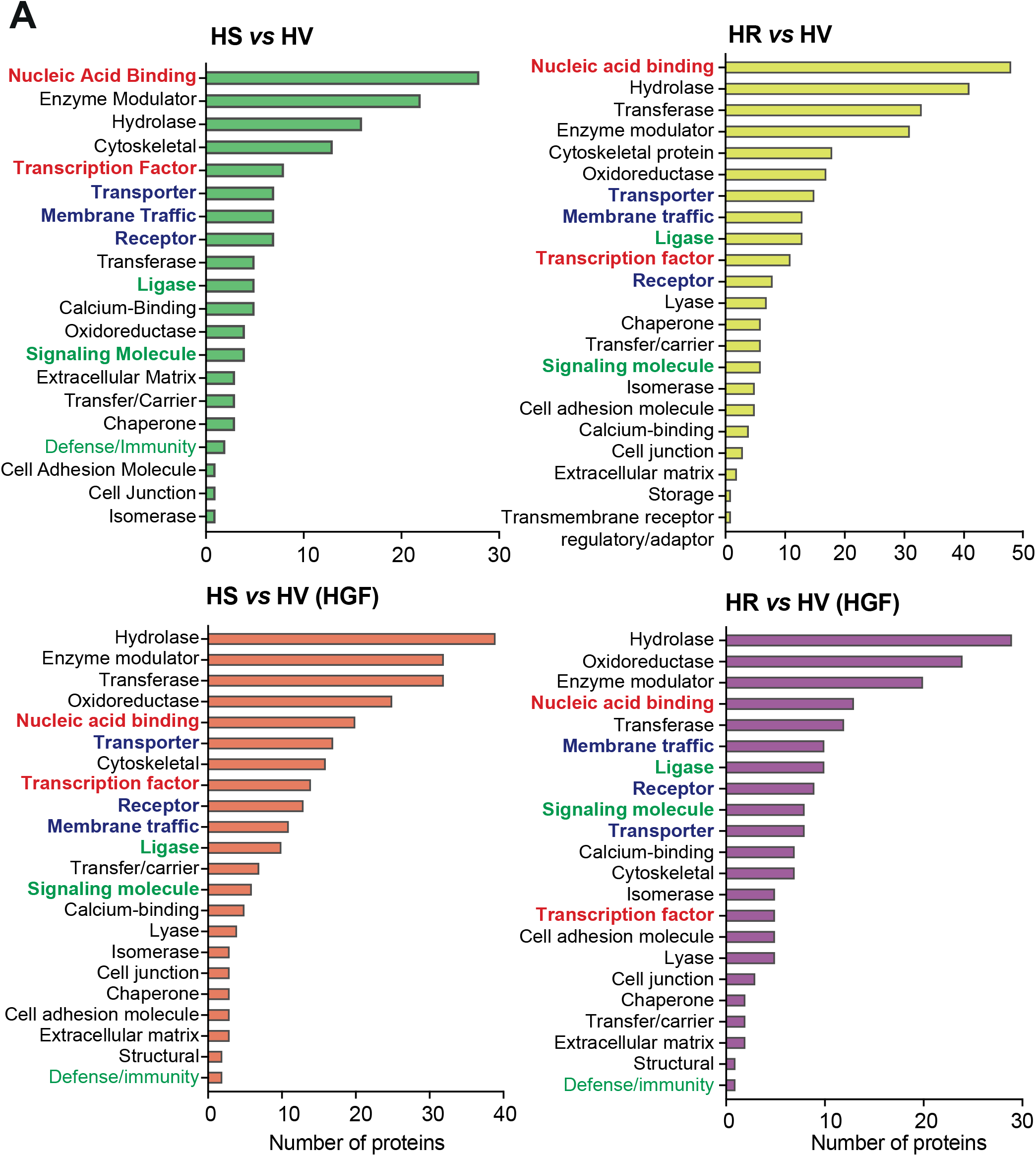
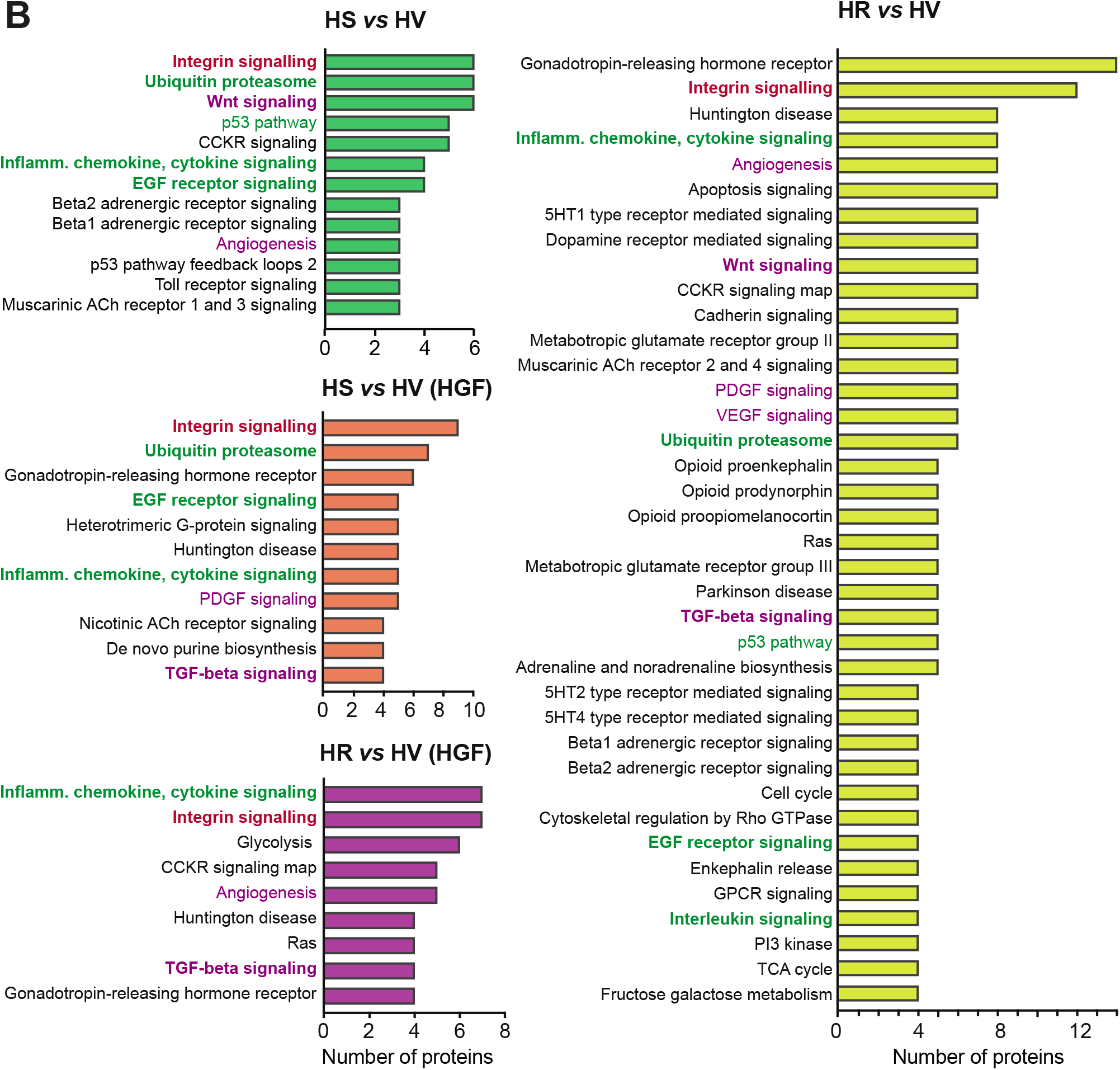
SOCS1 modulates diverse classes of proteins in multiple signaling pathways. Proteins that were significantly modulated (both downregulated and upregulated) by SOCS1 at steady state (HS *vs* HV) or after HGF stimulation (HS *vs* HV (HGF)), and by SOCS1R015K at steady state (HR *vs* HV) or after HGF stimulation (HR *vs* HV(HGF)) were analyzed using the PANTHER database. The protein classes (A) and the cellular signaling pathways (B) that contained the highest numbers of modulated proteins are shown. The protein classes and pathways shared in multiple comparisons are indicated by the same colored font.

SOCS1 is a well-known regulator of the JAK-STAT and RTK signaling pathways and is a substrate adaptor for protein ubiquitination [16,42]. Even though the inflammatory chemokine and cytokine, ubiquitin-proteasome and EGF receptor signaling are among the most prominent pathways modulated by SOCS1 at steady state and after HGF stimulation (Fig. 2B; HS *vs* HV and HS *vs* HV (HGF)), their rank order was altered following HGF stimulation. Disruption of the phospho-Tyr binding FLVRS motif within the SH2 domain (SOCS1R105K) did not affect the regulation of these three pathways at steady state (HR *vs* HV) but abrogated the ubiquitin-proteasome and the EGF receptor signaling pathways in HGF-stimulated cells (HR *vs* HV (HGF)). The members of these modulated pathways are identified in Supplementary Tables S3A-S3D.

Intriguingly, the integrin signaling pathway proteins ranked the top among those modulated by SOCS1 at steady state and after HGF stimulation, and the loss of phospho-tyrosine binding motif doubled the number of proteins within this pathway that was reduced by HGF stimulation (Fig. 2B). Other pathways prominently altered by SOCS1 in p-Tyr-independent manner are the Wnt signaling and p53 pathways, which were not represented in HGF-stimulated cells. Strikingly, the number of pathways altered by SOCS1 profoundly increased when the p-Tyr binding pocket was disrupted (HR vs HV). However, this increase in diversity and number of proteins was reversed following HGF stimulation. Conversely, the TGFβ signaling pathway, which was not observed in Hepa-SOCS1 cells, emerged after HGF stimulation and in Hepa-SOCS1R105K cells at steady state and after HGF stimulation (Fig. 2B). Lastly, SOCS1 and the SH2 domain mutant altered a number of proteins of the adrenergic receptor and other GPCR signaling pathways (Fig. 2B; Supplementary Tables S3C, S3D).

### Loss of p-Tyr binding and HGF signaling profoundly alters SOCS1-modulated proteins

The signaling pathways altered by SOCS1R105K at steady state were more numerous and substantially differed compared to those altered by WT SOCS1 (Fig. 2B). Therefore, we determined how the loss of p-Tyr binding motif and activation of the MET tyrosine kinase influenced SOCS1-dependent protein modulation through pairwise comparisons of the four conditions through Venn diagrams and heatmaps. Strikingly, SOCS1R105K modulated more proteins than WT SOCS1 at steady state and there was minimal overlap between the two groups (Fig. 3A, Venn diagram). The ten proteins shared between the two cells showed an inverse modulation, with proteins downregulated by SOCS1 becoming upregulated in cells expressing SOCS1R105K, and vice versa (Fig. 3A, heatmap). These observations suggest that the p-Tyr binding pocket may serve to maintain the specificity of SOCS1-dependent protein modulation. The number of proteins modulated by WT SOCS1 after HGF stimulation more than doubled compared to proteins modulated at steady state, and these two groups also showed very little overlap (Fig. 3B). Among the nine shared proteins between these two groups, only ACIN1 showed downregulation in both and the rest showed an inverse relationship. These findings suggest that the MET RTK activity stimulated by HGF may generate numerous new substrates that divert the action of SOCS1 from their steady state substrates.

**Fig. 3.**
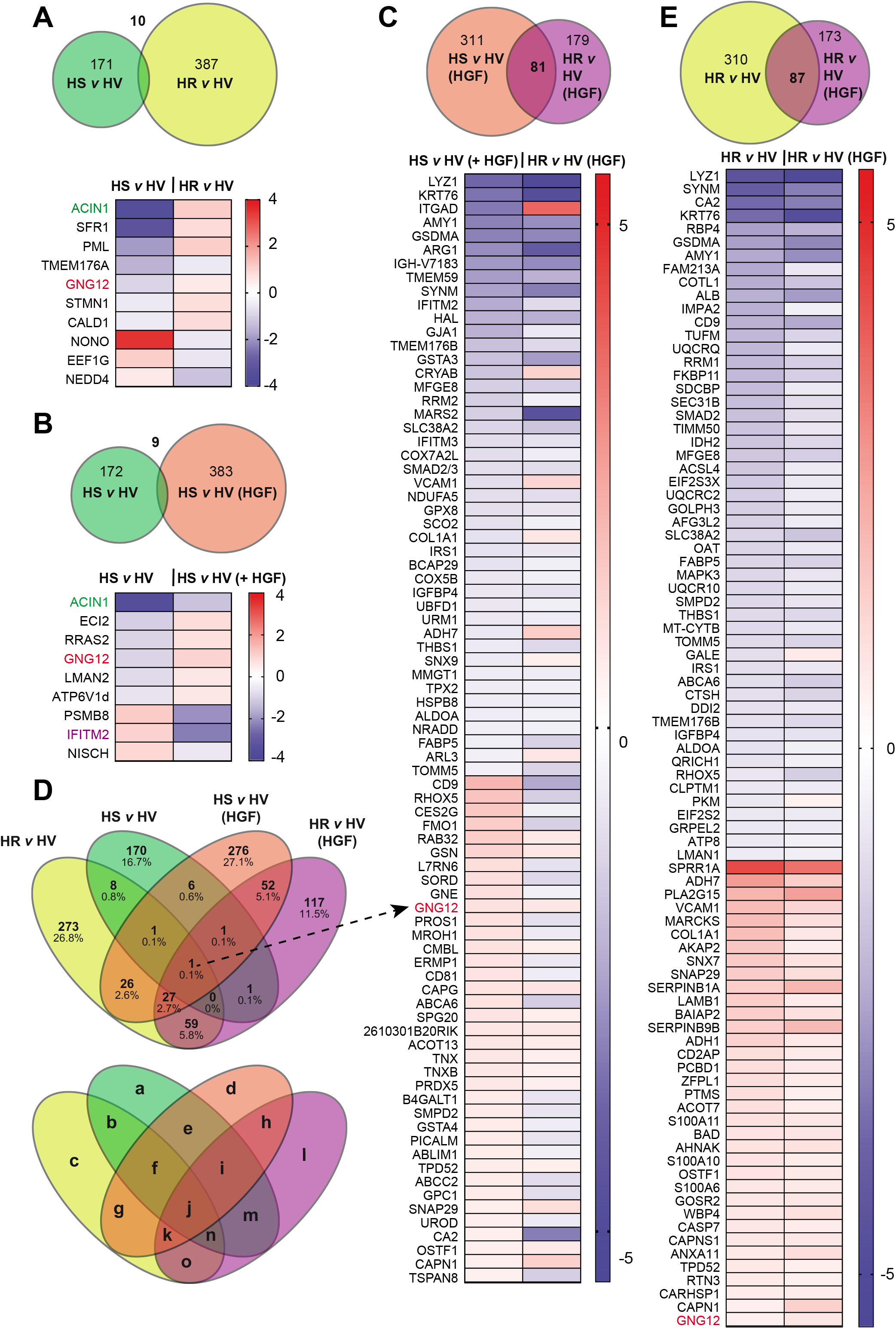
Loss of p-Tyr binding and HGF signaling profoundly alter SOCS1-modulated proteins. Proteins modulated by WT SOCS1 or the SH2 domain mutant (SOCS1R105K) in cells at steady state or after HGF stimulation were analyzed by pairwise comparisons: (A) Venn diagram of proteins modulated by WT SOCS1 and SOCS1R015K at steady state. The shared proteins and their down- or up-regulation are indicated in heatmaps. (B) Proteins modulated by WT SOCS1 at steady state and after HGF stimulation. (C) Proteins modulated by WT SOCS1 and SOCS1R015K after HGF stimulation. (D) Proteins modulated by SOCS1R015K at steady state and after HGF stimulation. (E) 4-way Venn diagram of proteins modulated in all four groups. The numbers are percentages are given in the upper panel, and the lower panel designates the various sets as letters. The list of proteins in each set are listed in Supplementary Table S4.

Disrupting the p-Tyr binding pocket of SOCS1 reduced the number of proteins modulated by WT SOCS1 following HGF stimulation (Fig. 3C; 311+81 vs 179+81). However, there was considerable overlap between the proteins modulated in these two groups, although more than one third of the 81 common proteins showed differential modulation in HGF-stimulated cells expressing WT SOCS1 or SOCS1R105K (Fig. 3C, heatmap: 29 out of 81). These observations suggest that the similarly affected proteins may represent p-Tyr independent modulation, whereas the differentially affected proteins likely represent p-Tyr dependent modulation. The former group (modulated in a p-Tyr independent manner) is represented in set ‘h’ of a 4-way Venn diagram as proteins shared only between these two groups (Fig. 3D, set ‘h’; 52 out of 81). The list of proteins in each set of the 4-way Venn diagram is given in Supplementary Table S4. There was also a considerable overlap between proteins modulated by SOCS1R105K at steady state and following HGF stimulation (Fig. 3D, set ‘k’, 27 out of 81). These observations suggest that proteins modulated by SOCS1R105K both at steady state and after HGF stimulation are not the steady state substrates of WT SOCS1 and reinforce the notion that these proteins are targeted when the ability of SOCS1 to interact with its regular partners is compromised. Consistent with this possibility, proteins modulated by SOCS1R015K at steady state and following HGF stimulation showed the maximal overlap and they were modulated in the same way under both conditions (Fig. 3E). Furthermore, GNG12 is the only protein modulated in all four comparisons. GNG12, which is induced following LPS stimulation [43], was downregulated in HS cells but upregulated in HR cells.

### SOCS1 modulates proteins of macromolecular processing complexes

Analysis of protein interaction networks using the STRING database revealed prominent clustering of proteins involved in macromolecular homeostasis among proteins downregulated or upregulated by WT SOCS1 at steady state (Fig. 4). The most prominent among these clusters are members of the E2 ubiquitin (Ub) conjugation enzymes (UBC) coded by several *Ube2* genes [44,45]. These proteins catalyze the second step in protein ubiquitination by transferring the activated Ub and Ub-like moieties to one of the numerous E3 Ub ligases. SOCS1 modulated several UBE2 proteins, upregulating UBE2D1, UBE2D2A, UBE2DB, UBE2D3, UBE2V1 and GM20431 (a predicted protein with UBC activity), and downregulated UBE2R2 and UBE2V2 (Fig. 4). All these UBE2 proteins are involved in transferring the Ub moiety and not the other Ub-like moieties such as SUMO, NEDD8 or ISG15 [44]. A related macromolecular assembly influenced by SOCS1 is the proteasome, which degrades ubiquitinated proteins into peptides, some of which are loaded on to MHC class-I molecules for antigen presentation [46,47]. The structure of the proteasome and its various segments, and their protein components are illustrated in the Supplementary Figure S2. SOCS1 downmodulated the core proteasome components PSMB1 and PSMB7, but upregulated the immunoproteasome core component PSMB8, which replaces PSMB5 [47] (Fig. 4, Supplementary Figure S2). Intriguingly, SOCS1 expressing cells also upregulated PSMF1, which functions as the physiological proteasome inhibitor PI31 [48]. These data suggest that SOCS1, in addition to serving as substrate adaptor for CUL5^SOCS1^ E3 Ub ligase, has the potential to modulate the ubiquitination and proteasome machineries that may have an impact on overall cellular protein homeostasis.

**Fig. 4.**
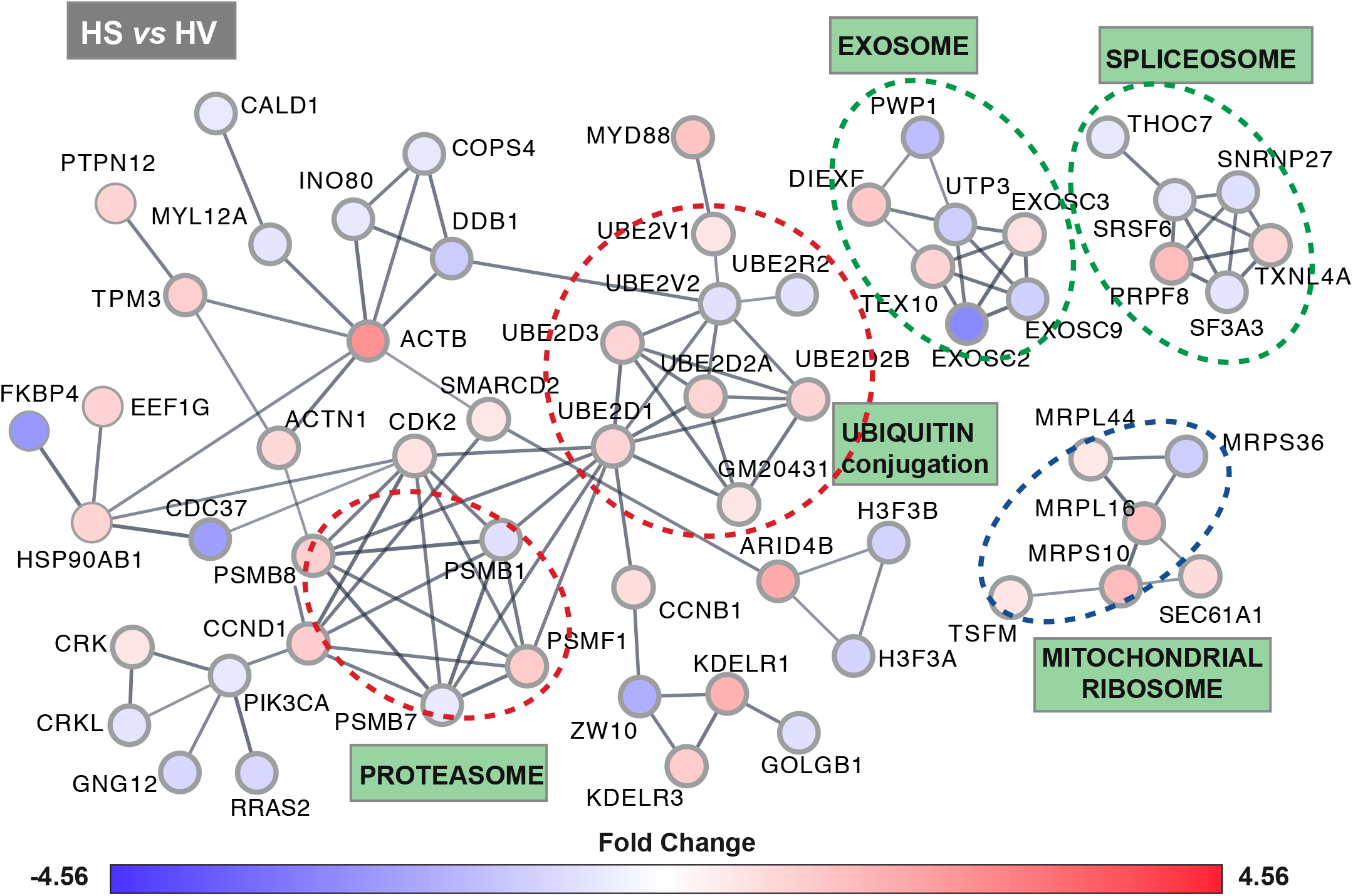
SOCS1 modulates proteins of macromolecule processing complexes. (A) Proteins that are significantly modulated by SOCS1 (both downregulated and upregulated) were analyzed using the STRING database to identify the cellular protein networks influenced by SOCS1. Enrichment of the SOCS1-modulated proteins within the ubiquitin conjugating enzymes, and components of the proteasome, the RNA spliceosome, the RNA exosome and the mitochondrial ribosome are encircled. Color of the protein circles reflects SOCS1-mediated fold change in expression as indicated by the heatmap. Note that SOCS1 upregulates certain components but downregulates others in the identified multi-protein complexes.

The protein interaction network analysis revealed clusters of SOCS1-modulated proteins that constitute the RNA spliceosome and RNA exosome, multi-protein complexes involved in RNA splicing and turnover, respectively (Fig. 4). The RNA spliceosome is a very complex machinery composed of protein chaperones that organize catalytic small nuclear RNAs (snRNA) that remove intronic sequences from primary transcripts, as they are produced, to generate functional mRNA and long non-coding RNAs (lncRNA) [49,50]. SOCS1-expressing cells at steady state downregulated SF3A3, SRSF6 and SNRNP27, and upregulated PRPF8 and TXNL4A, which are implicated in RNA splicing. THOC7, involved in transcription termination and RNA transport, was also downregulated in HS cells. The RNA exosome is a ribonuclease complex that functions in nuclear and cytoplasmic RNA surveillance pathways that ensure accurate processing of RNA precursors, degrade inherently unstable and aberrantly processed transcripts, and participate in mRNA turnover [51–53]. The RNA exosome is very similar in organization and function to the proteasome, composed of 3’ to 5’ exonucleases in the inner core and outer regulatory subunits, which functions in concert with diverse co-factors that recognize the RNA structure/sequence [52]. SOCS1 expressing cells at steady state showed downregulation of EXOSC2 and EXOSC9, and upregulation of EXOSC3 of the RNA exosome core (Fig. 4). Their interactome contained proteins involved in rRNA processing (downregulated: UTP3 and PWP1; upregulated: DIEXF and TEX10).

The other prominent SOCS1-modulated interactome contained protein constituents of the mitochondrial ribosome, which is essential for translating all mitochondrial and many nuclear mRNAs [54,55]. SOCS1 differentially modulated the mitoribosome small subunit proteins MRPS36 (down) and MRPS10 (up), and upregulated the large subunit proteins MRPL16 and MRPL44, as well as TSFM (Fig. 4), a mitochondrial translation elongation factor [56]. This interactome also included SEC61A1, a component of the SEC61 translocon complex regulating protein translocation from ribosomes into endoplasmic reticulum [57,58].

Network analysis of protein modulated in other pairwise comparisons HS *vs* HV (HGF), HR *vs* HV and HR *vs* HV (HGF) are shown in Supplementary figures S3A to S3C, respectively. Notable protein groups in the HR *vs* HV group included several mitochondrial proteins. In the HS *vs* HV (HGF) group, the UBE2 group of proteins showed an inverse trend compared to the HS *vs* HV group (Fig. 4, Supplementary figures S3A). Therefore, we investigated modulation of the Ub-proteasome pathway proteins in more detail.

### SH2 domain mutation and RTK signaling profoundly alters the UBE2 enzymes and proteasome components modulated by SOCS1

To determine whether the modulation of UBE2 enzymes and proteasome components by SOCS1 is dependent on its SH2 domain and whether it is influenced by RTK signaling, we compared the expression of all members of these two protein classes that were detected by the SILAC mass tag in the total proteome of HS cells compared to HV cells (Fig. 5). SOCS1 significantly modulated the expression of UBE2D, UBE2R2, UBE2V1 and UBE2V2, which were reversed by the SH2 domain mutation (Fig. 5A). Even though SOCS1 strongly modulated several other UBE2 family members (e.g., down: UBE2F, UBE2M; up: UBE2B, UBE2I), these changes were not significant (FDR >0.05). On the other hand, SOCS1R105K expression altered the expression of other UBE2 proteins namely, UBE2G1, UBE2H and UBE2L3. Activation of the MET RTK also caused significant changes in the expression UBE2 proteins in cells expressing WT SOCS1: UBE2V1 upregulated at steady state was downregulated in HGF-stimulated cells; the latter also showed significant downregulation of UBE2C, UBE2E, UBE2I, UBE2N and UBE2S. Most of these HGF-induced changes in WT SOCS1 expressing cells were reversed in SOCS1R015K expressing cells.

**Fig. 5.**
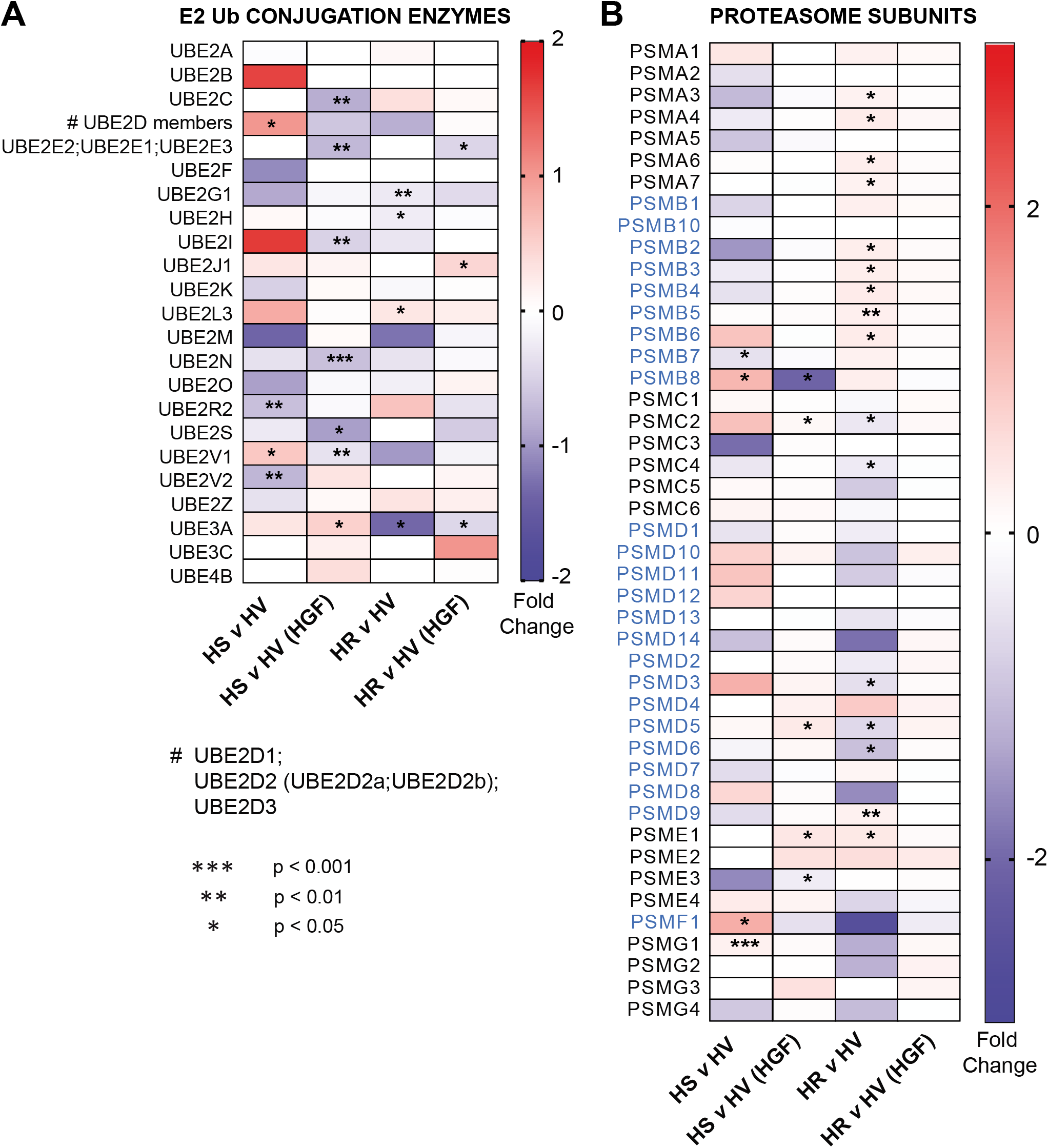
SH2 domain mutation and RTK signaling profoundly alters the UBE2 enzymes and proteasome components modulated by SOCS1. (A) Heatmap of ubiquitin conjugation enzymes modulated by SOCS1 or SOCS1R105K at steady state and after HGF stimulation. The four classes of UBE2 enzymes are color coded. ^*#*^UBE2D includes four members: UBE2D1, UBE2D2 (UBE2D2A, UBE2D2B) and U UBE2D3. A subset of Ub conjugation enzymes UBE3A, UBE3C and UBE4B are also shown. (B) Heatmap of proteasome components modulated by SOCS1 or SOCS1R105K at steady state and after HGF stimulation. The protein components of the various segments of the proteasomes namely the outer core (PSMA), inner core (PSMB), base (PSMC) and lid (PSMD) of the regulatory subunits, activator (PSME) and inhibitors of the proteasome (PSMF), and the proteasome assembly chaperones (PSMG) are identified by different colors (see Supplementary Fig. S2 for details). Significant changes are indicated by asterisks: *p<0.05, **p<0.01, ***p<0.001.

Most components of the proteasome showed some changes in Hepa-SOCS cells at steady state, but only a few of them namely, PSMB7, PSMB8, PSMF1 and PSMG1, were significantly altered by SOCS1 (Fig. 5B). Strikingly, mutation of the SH2 domain caused widespread changes in the proteasome composition, significantly upregulating many proteins of the outer (PSMA3, 4, 6, 7) and inner (PSMB2, 3, 4, 5, 6) rings of the 20S core proteasome, while downregulating some proteins of the 19S regulatory particle (PSMD3, 5, 6) (Fig. 5B, Supplementary Fig. S2). HGF stimulation caused much fewer changes in SOCS1-modualted proteasome components, the most prominent being downregulation of PSMB8, which was elevated at steady state. Strikingly, HGF stimulation reversed all changes observed at steady state in cells expressing the mutant SOCS1. These observations indicate that SOCS1 has the potential to profoundly alter cellular protein homeostasis by modulating the ubiquitination and proteasomal degradation machineries.

### Modulation of UBE2D proteins by SOCS1 occurs at the post-transcriptional level

The UBE2D proteins include the Ub conjugating enzymes UbcH5A, B, C and D that were previously known as Ubc4/5 [45]. A recent study identified UBE2D enzymes as interacting partners of the phosphorylated E3 ligase CBL, which is implicated in the ubiquitination of EGFR and MET receptors [59]. Moreover, Ubc5 was previously implicated in the ubiquitination functions of the von Hippel-Lindau protein, the prototype of SOCS box containing Ub ligase complexes [22,60]. As deregulated expression of CBL, RTKs, VHL and SOCS1 are associated with malignant growth, and SOCS1 has been reported to promote MET ubiquitination [9], we first examined the relationship between the expression of SOCS1, CBL, MET and UBE2D in TCGA-LIHC dataset. *SOCS1* expression correlated positively with that of *UBE2D1* and *UBE2D2* (Fig. 6), similarly to the results on protein expression in HS cells (Fig. 5A), but not with *UBE2D3*. The expression of *SOCS1* and *CBL* negatively correlated with that of *MET*, and expression of *UBE2D1* and *UBE2D2* showed a similar negative correlation with that of MET (Fig. 6).

**Fig. 6.**
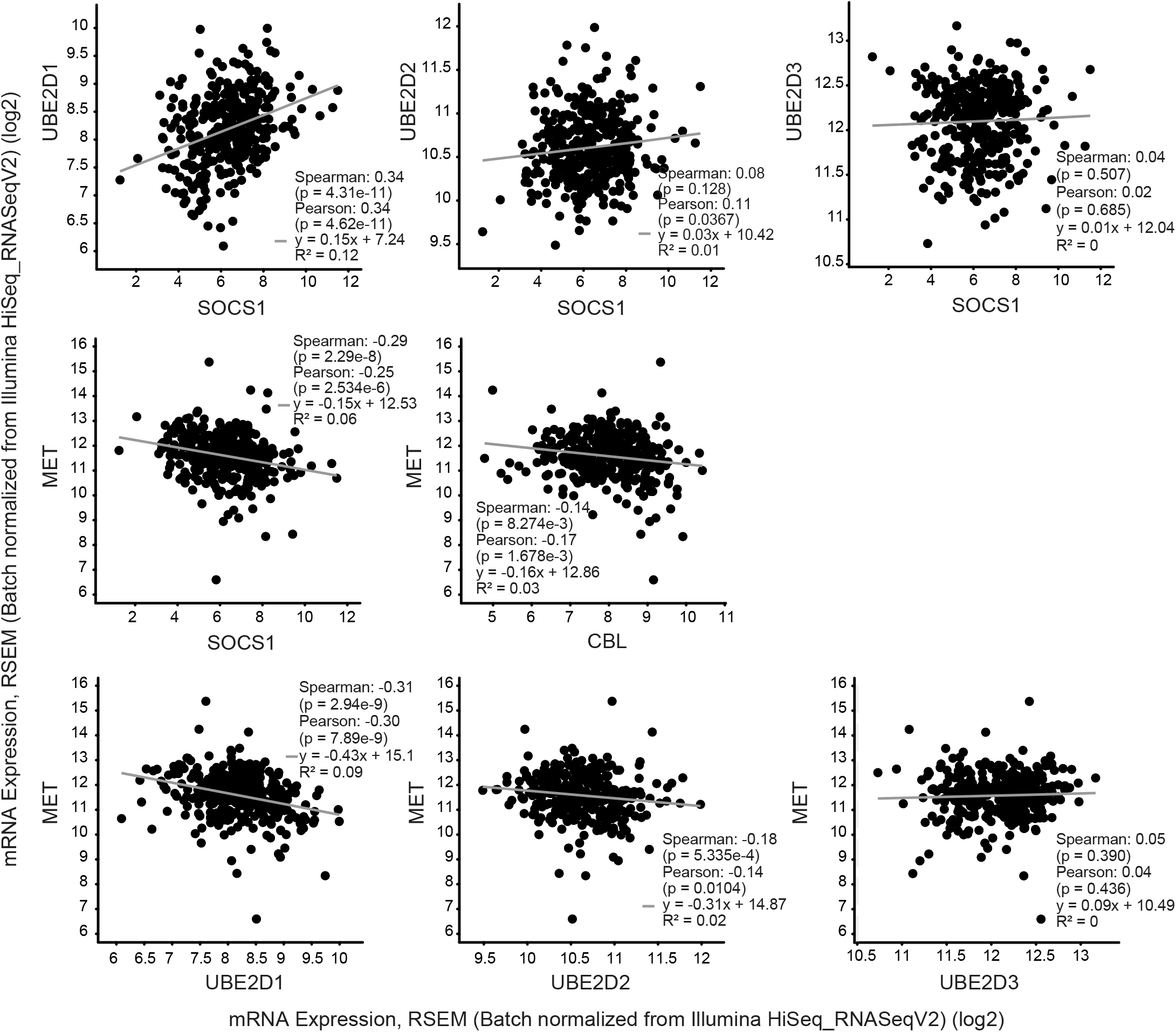
Correlation between the expression levels of *SOCS1* and *UBE2D2, CBL* and *MET* in the TCGA-liver hepatocellular carcinoma dataset.

The above results suggested that SOCS1-dependent regulation of UBE2D in HS cells may occur at the transcriptional level, and that UBE2D, as a component of both CBL- and SOCS-containing E3 ubiquitination ligases, could downmodulate MET expression. Therefore, we examined the transcript levels of *Ube2d* genes in Hepa cells. Contrary to our expectation, the transcript levels of *Ube2d1* and *Ube2d3* expression levels were significantly reduced in HS cells at steady state, whereas there was no appreciable change in *Ube2d2* mRNA levels (Fig. 7A). HGF stimulation caused strong transcriptional upregulation of *Ube2d2* that was further augmented by SOCS1 at 8h after HGF stimulation that stabilized over 16-24h. On the other hand, HGF stimulation decreased the expression of *Ube2d1* and *Ube2d3*. These results indicated that the increased expression of UBE2D proteins in HS cells at steady state observed in SILAC experiments likely resulted from changes in protein levels rather than from increased gene expression.

**Fig. 7.**
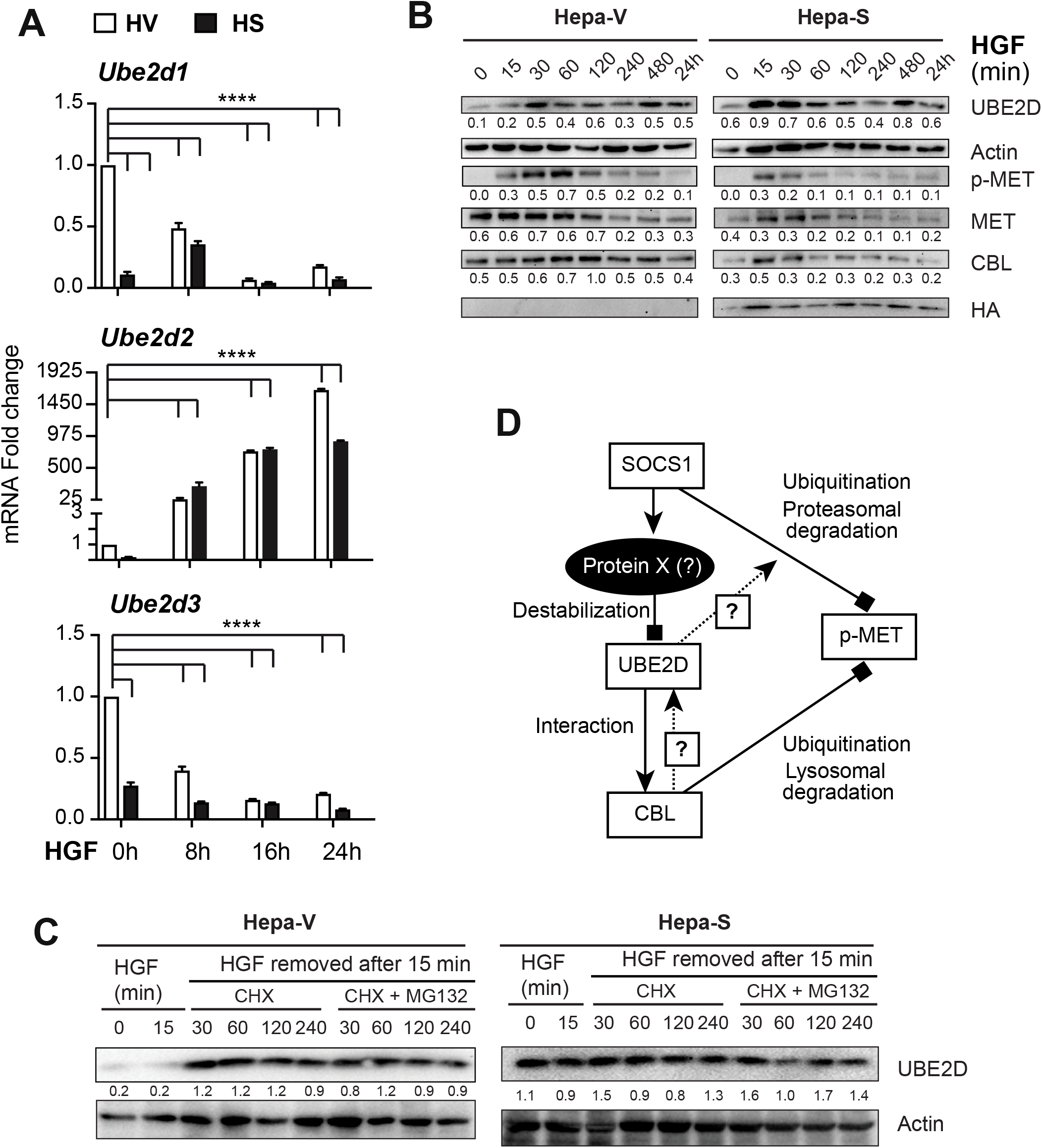
Effect of SOCS1 on the expression of UBE2D, CBL and MET proteins after HGF stimulation. (A) Expression of *UBE2D* mRNA in HV and HS cells at steady state and at different time pints after HGF stimulation. Pooled data from three experiments are shown. (B) Modulation of the protein levels of UBE2D, phosphor-MET, MET and CBL in HV and HS cells after HGF stimulation. (C) Stabilization of UBE2D in HV cells in the presence of cycloheximide. For (B) and (C), representative data from two experiments are shown, and numbers below the protein bands indicate the ratio compared to actin expression at the corresponding time point. (D) Proposed model implicating an unknown protein regulated by SOCS1 in controlling the expression of UBE2D and the role of UBE2D in modulating the turnover of activated MET receptor.

### Downregulation of MET by SOCS1 following HGF stimulation correlates with downregulation of CBL and upregulation of UBE2D

An antibody that recognizes all isoforms namely, UBE2D1, UBE2D2 and UBE2D3 was used to evaluate the expression of UBE2D proteins. Hepa-SOCS1 showed discernibly elevated basal levels of UBE2D compared to control cells (Fig. 7B; time 0), supporting the prediction that elevated UBE2D levels in HS cells results from post-transcriptional regulation. In addition, these results indicated that UBE2D protein levels are regulated at steady state and that SOCS1 deregulates this control. Both control and SOCS1 expressing cells markedly upregulated UBE2D levels after 3 hours of HGF stimulation (Fig. 7B), indicating that the UBE2D protein levels are modulated by RTK signaling. We confirmed HGF-induced Tyr phosphorylation of MET and observed downmodulation of both phospho-MET and total-MET by SOCS1 (Fig. 7B) as previously reported [9]. We also observed that CBL, which binds to phosphorylated MET and targets it for lysosomal degradation [61], was also reduced by SOCS1 following HGF stimulation (Fig. 7B). Downmodulation of both MET and CBL in HGF-stimulated cells expressing WT SOCS1 correlated with elevated levels of UBE2D at steady state and at the early time points of HGF stimulation. Collectively, these findings suggest that upregulation of UBE2D in HS cells is coupled to SOCS1-mediated downmodulation of MET along with CBL, which is recruited to the activated MET receptor.

### Upregulation of UBE2D by SOCS1 does not require new protein synthesis

To determine if increased UBE2D levels in Hepa-S cells required new protein synthesis, and to determine whether its decrease in Hepa-V cells is mediated by proteasomes, we exposed the cells to HGF for 15 min, washed them and incubated in the presence of the protein translation inhibitor cycloheximide (CHX) either alone or along with the proteasome inhibitor MG132. We observed that not only CHX did not reduce UBE2D levels in HS cells but also upregulated UBE2D in HV cells. Addition of MG132 did not cause any further increase in UBE2D levels already upregulated by CHX in HV cells (Fig. 7C). These findings suggest that UBE2D levels in Hepa cells are regulated by an unknown protein, which requires *de novo* protein synthesis and is subject to SOCS1-mediated regulation (Fig. 7D). This would explain the stabilization of UBE2D in CHX-treated HV cells and in HS cells at steady state.

## Discussion

In this study, we show that (i) SOCS1 can potentially modulate the expression of hundreds of proteins in hepatocytes that impact on multiple signaling pathways, (ii) this profile is profoundly altered by the R105K mutation within the SH2 domain of SOCS1 or by strong tyrosine kinase signaling, (iii) a quarter of SOCS1-modulated proteins in HGF-stimulated cells are not influenced by the loss of p-Tyr binding capacity of SOCS1, and (iv) among the proteins modulated by SOCS1, clusters of proteins that constitute large multiprotein complexes such as the proteasome, the RNA spliceosome, the RNA exosome and the mitoribosome predominate. Besides, our study finds that SOCS1 modulates several members of the UBC enzymes involved in protein ubiquitination, among which UBE2D (UBC4/5) appears to be stabilized by SOCS1 indirectly through another unidentified SOCS1 regulated proteins. As SOCS1 is induced by a diverse array of cell stimuli such as interferons, interleukins, growth factors, chemokines, hormones and endogenous glucocorticoids, and microbial products such as LPS and fMLP either directly or indirectly [62–64], we propose that SOCS1, in addition to attenuating the signaling pathways activated by these stimuli may also help the cells to reset cellular protein homeostasis through modulating RNA processing and protein ubiquitination and degradation.

Following the first report demonstrating the substrate adaptor function of SOCS1 for protein ubiquitination of VAV1, many cellular proteins such as JAK2, IRS1, IRS2, FAK and EPOR have been shown to undergo SOCS1-dependent ubiquitination and proteasomal degradation [16,21]. Almost all these studies used overexpression of SOCS1 as well as the target substrates in model cell systems to document SOCS1-dependent modulation. Very few studies have examined the effect of an endogenous substrate in cells stably expressing SOCS1, or the effect of SOCS1 loss on endogenous substrate [9,65]. Recently, a shotgun mass spectrometry was used to identify SOCS1-interacting proteins following immunoprecipitation of overexpressed SOCS1 in the acute myeloid leukemia (AML) cell line KG-1 [17]. This study, aimed at identifying protein(s) that downmodulate SOCS1 expression in AML at the posttranscriptional level, identified 87 SOCS1-interacting proteins. Even though 26 of these proteins were modulated in SOCS1-dependent manner in our study, only two proteins namely, mitochondrial single strand DNA binding protein (SSBP1) and CT-10 regulator of kinase (CRK)-like protein (CRKL) showed statistically significant downmodulation. Both proteins are implicated in malignant growth of multiple cancers. SSBP1 is implicated in protecting cancer cells from proteotoxic stresses, promoting mitochondrial replication and conferring radio-resistance [66–68]. CRKL, containing SH2 and SH3 domains, can impact on myriad of cellular signaling pathways and is implicated in neoplastic growth [69–71]. A recent report implicated CRL in promoting alternate splicing of cancer related genes [72]. These reports raise the possibility that SOCS1 may mediate its tumor suppressor functions at least partly via downmodulating SSBP1 and CRKL.

We used SILAC-based proteomics to maximize the detection of all potential targets of SOCS1-mediated protein modulation. Even though our initial focus was on protein downmodulation mediated by the ubiquitination function of SOCS1, many proteins were also found to be upregulated in HS cells. Such increase in protein expression could result from SOCS1-induced changes in the expression of transcriptional factors as well as regulators of mRNA stability and translation might contribute to protein upregulation. Indeed, nucleic acid binding proteins and transcription factors were among the prominent protein classes altered by SOCS1 (Fig. 2A). Besides, modulation of the protein ubiquitination process could also contribute to the upregulation of certain proteins, as exemplified by increased expression of the UBC enzyme UBE2D. Our findings suggest that the increased UBE2D expression in HS cells results from its stabilization, presumably resulting from SOCS1-mdeiated loss of an unknown protein(s) that keeps UBE2D levels low in control cells.

It is possible that CBL itself may be a regulator of UBE2D levels. CBL is an E3 Ub ligase that promotes ubiquitination and lysosomal sorting and degradation of RTKs such as EGFR and MET [61,73]. Proteasomes were also implicated in the regulation of MET and we have implicated SOCS1 in this process [42,74]. However, it remains unclear whether these two pathways are regulated, and if so, how the preference for one pathway over the other is achieved. CBL itself is regulated by complex mechanisms involving Ub-dependent lysosomal degradation of RTK-associated and proteasomal degradation of cytosolic CBL [75]. Even though UbcH7 (UBE2L3) was initially implicated in CBL-mediated EGFR ubiquitination, a later report showed that UBC4/5 (UBE2D members) were involved [44,76,77]. The Lipkowitz lab recently showed that in addition to UBE2D also promotes auto-ubiquitination of CBL [59]. This study also showed that all these UBE2 and UBE2W promote auto-ubiquitination of CBL. Whether UBE2D is also involved in SOCS1-dependent ubiquitination of MET remains to addressed (Fig. 7D, dashed line). As SOCS1 reduces the expression of CBL upon MET activation, it is likely that SOCS1-dependent downmodulation of MET also promotes degradation of the MET-associated CBL and this may lead to stabilization of UBE2D proteins (Fig. 7D, dashed line). Clearly further studies are needed to decipher the consequences of UBE2D-CBL interaction, and how this is modulated by SOCS1-dependent RTK regulation.

Given that SOCS1 is ubiquitously induced by cytokines, growth factors and TLR stimulation, our findings suggest a complex role of SOCS1 in maintaining cellular functions in hepatocytes that may also be applicable to other cell types. As SOCS1 expression is tightly regulated, a potential limitation our study using cell lines stably expressing SOCS1 could be that the cell lines could become adapted over time. Nonetheless, our study suggest that the SOCS1-dependent regulation of cellular macromolecular complexes could probably be invoked under conditions of increased SOCS1 expression induced by cellular activation signals. SOCS1-deficient mice are born normal but succumb to widespread systemic inflammation affecting the liver and other vital organs [78,79], indicating the indispensable role of SOCS1 in maintaining cellular homeostasis. The lethal phenotype of SOCS1 is reversed by simultaneous ablation of the *Ifng* gene. IFNγ is a key modulator of the proteasome, which regulates protein turnover and also generates peptides for presentation by MHC class-I pathway to CD8^+^ cytotoxic T lymphocytes [80,81]. As CD8^+^ T cells are key mediators of autoimmune diseases and antitumor immunity [82,83], it is possible that the loss of SOCS1-dependent regulation of cellular macromolecular machineries may alter the profile of MHC-I bound peptides, and this may contribute to autoinflammatory responses. By the same token, repression of the *SOCS1* gene by epigenetic mechanisms in cancer cells may alter the Ub-proteasome pathway and compromise the development of antitumor immune responses. In this context, it is noteworthy that modulating the protein turnover in cancer cells has been shown to improve tumor antigen presentation [84]. Whereas SOCS1 has been studied extensively for its role in controlling cytokine and growth factor responses, our findings suggest that its potential functions in modulating cellular protein turnover and other macromolecular complexes need further exploration.

## Abbreviations

HCC: hepatocellular carcinoma
Hepa: Hepa 1-6 cell line
Hepa-R: Hepa 1-6 cells expressing SOCS1R105K mutant
Hepa-S: Hepa 1-6 cells expressing wildtype SOCS1
Hepa-V: Hepa 1-6 cells expressing control vector
HGF: hepatocyte growth factor
RTK: receptor tyrosine kinase
SILAC: Stable Isotopic Labelling of Amino acids in Cell culture
SOCS: suppressor of cytokine signaling.

## Availability of data and material

The mass spectrometry proteomics data have been deposited to the ProteomeXchange Consortium via the PRIDE partner repository with the dataset identifier PXD019135.

## Acknowledgements

The Canadian Institutes of Health Research (Project grant #PJT-153174 to SI) provided funding support to this project. MR was supported by the ‘Abdenour-Nabid MD’ graduate fellowship from the Faculty of Medicine, Université de Sherbrooke. The authors thank Ms. Catherine Allard (Statistician, CRCHUS) for writing the R package for determining the fold change and volcano plot analysis.

## Author Contribution

M.A.S. performed most of the experiments and data analysis; A.S. and M.C. performed the Western blot experiments; D.L. and F.M.B. performed the mass spectrometry and data collection; R.S. generated the cell lines; M.A.S. F.M.B. and S.I. designed the study, interpreted the data and wrote the manuscript.

## Notes

### Competing Interest Statement

The authors have declared no competing interest.

