## Supplementary Figures for "SILAC proteomics implicates the ubiquitin conjugating enzyme UBE2D in SOCS1-mediated downmodulation of the MET receptor in hepatocytes"

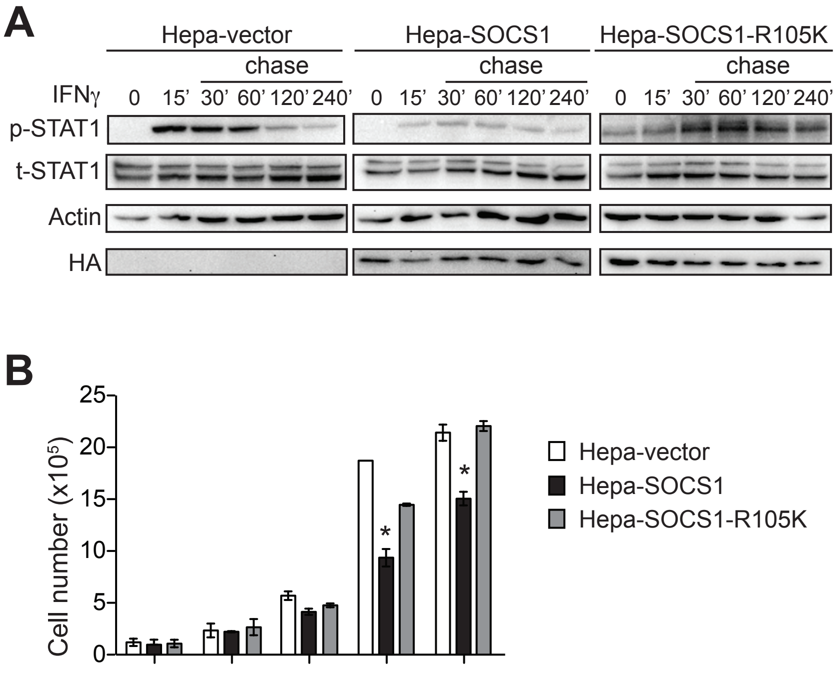


**Supplementary Fig. S1. Validation of the cell lines.** (A) Hepa1-6 cells expressing control vector (Hepa-Vector, HV), wild type SOCS1 (Hepa-SOCS1, HS) or the R105K mutant (Hepa-SOCS1R105K, HR) were incubated in DMEM containing 0.5% FBS (starving medium) over night and then exposed to IFNγ (20 ng/ml) in serum-free medium. Unstimulated (0') and IFNγ-stimulated cells were lysed immediately at 15 min (15'). Another set of stimulated cells were washed to remove IFNγ and incubated in fresh starving medium for 30', 60', 120' and 240' (chase). Cell lysates were probed for phospho-STAT1 (pSTAT1), total STAT1 (t-STAT1), actin or the HA tag of SOCS1 constructs. (B) 5 $x10^4$ HV, HS and HR cells were plated in 60 mm Petri dish in triplicates and cell counts were recorded at the indicated days. Media was replenished after 2 days, and cells were split into two 100 mm dishes after 4 days. Representative data from two independent experiments are shown for (A) and (B).

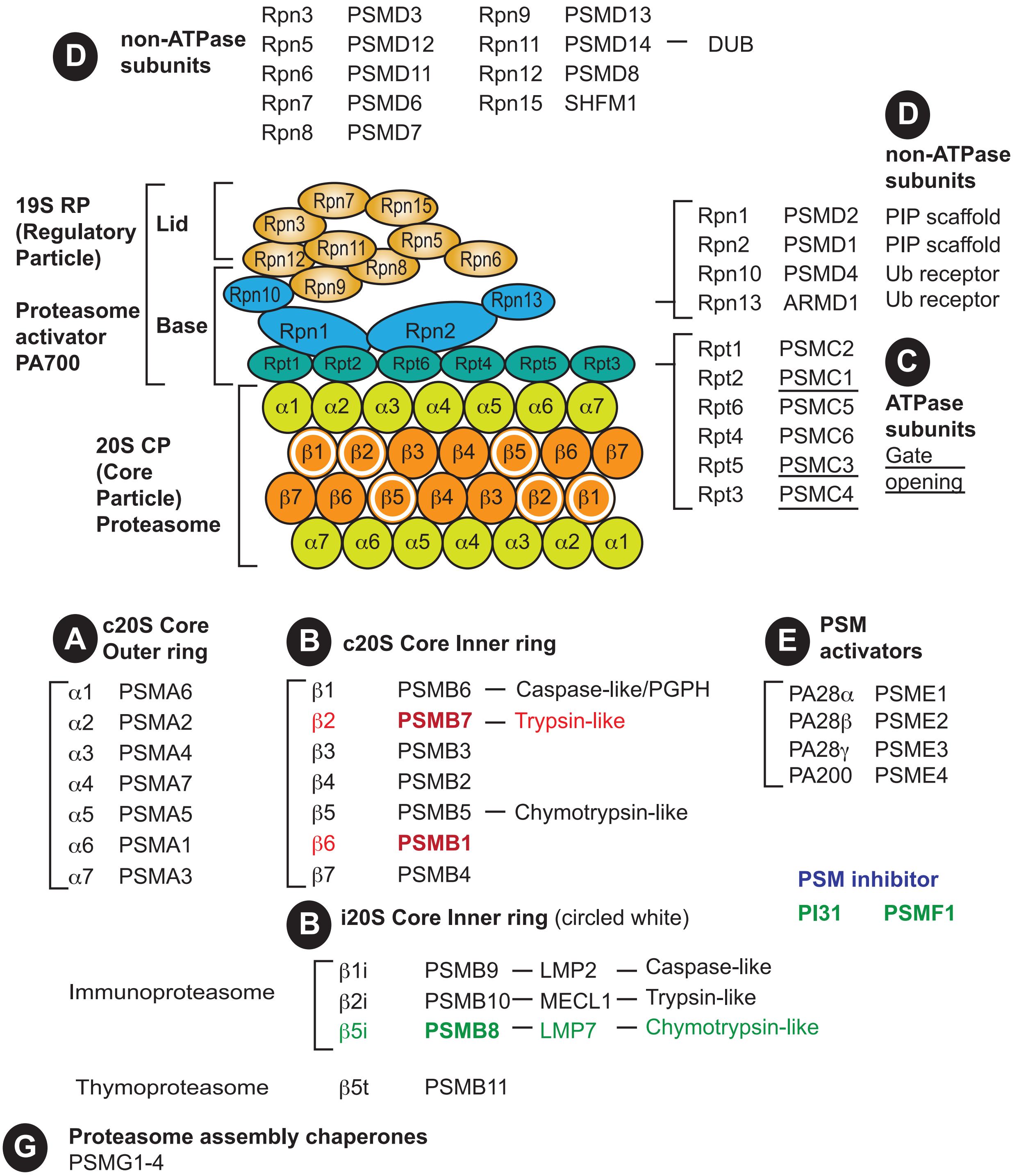


**Supplementary Fig. S2. Composition of the proteasome (PSM) and its regulators.** The protein constituents of the inner and outer rings of the 20S core particle, and the base and lid of the 19S regulatory subunits are illustrated. The inner ring subunits of PSM altered in immuno-PSM and thymo-PSM are indicated by white circles. Both traditional names and the HUGO nomenclature (PSM-A, B, C, D) are given. Changes in the compositions of the 20S PSM in immuno and thymo- PSM, as well as the PSM activators and inhibitors (PSM-E, F), and the PSM assembly chaperones (PSMG), are indicated: Red and green colors indicate up- and down- modulated proteins in HS *vs* HV cells. ***Abbreviations:*** **PIP**, Proteasome interacting protein; **PGPH**, peptidylglutamyl-peptide hydrolyzing; **PSM**, proteasome; **Rpn**, Regulatory particle, non-ATPase; **Rpt**, Regulatory particle, triple-ATPase.

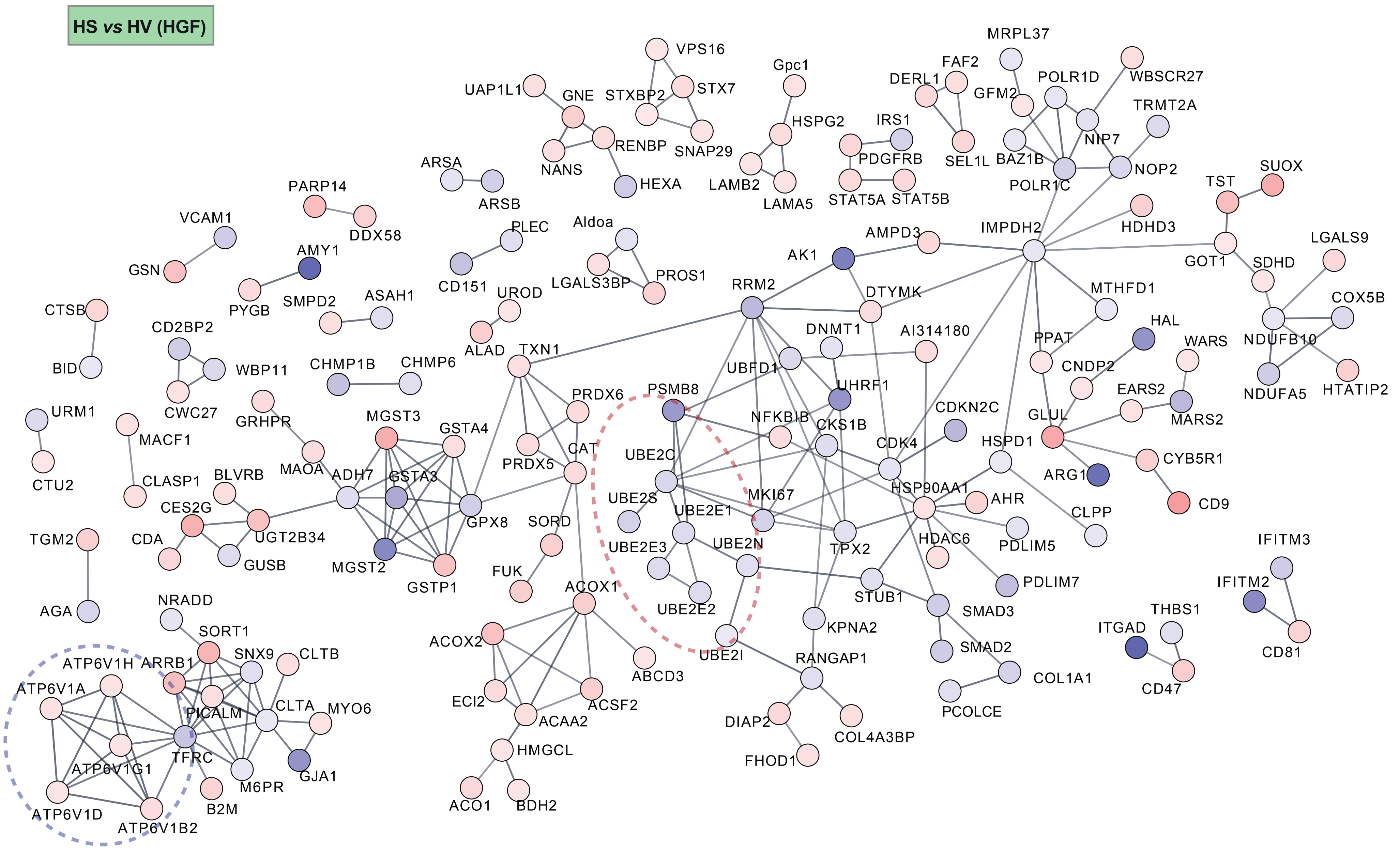


**Supplementary Fig. S3A. Network analysis of proteins modulated in HS cells following HGF stimulation.** UBC enzymes are circled in red and vascular ATPase, which regulate organelle acidification required for diverse protein procssing, in blue.

Fold Change

-4.07 4.07

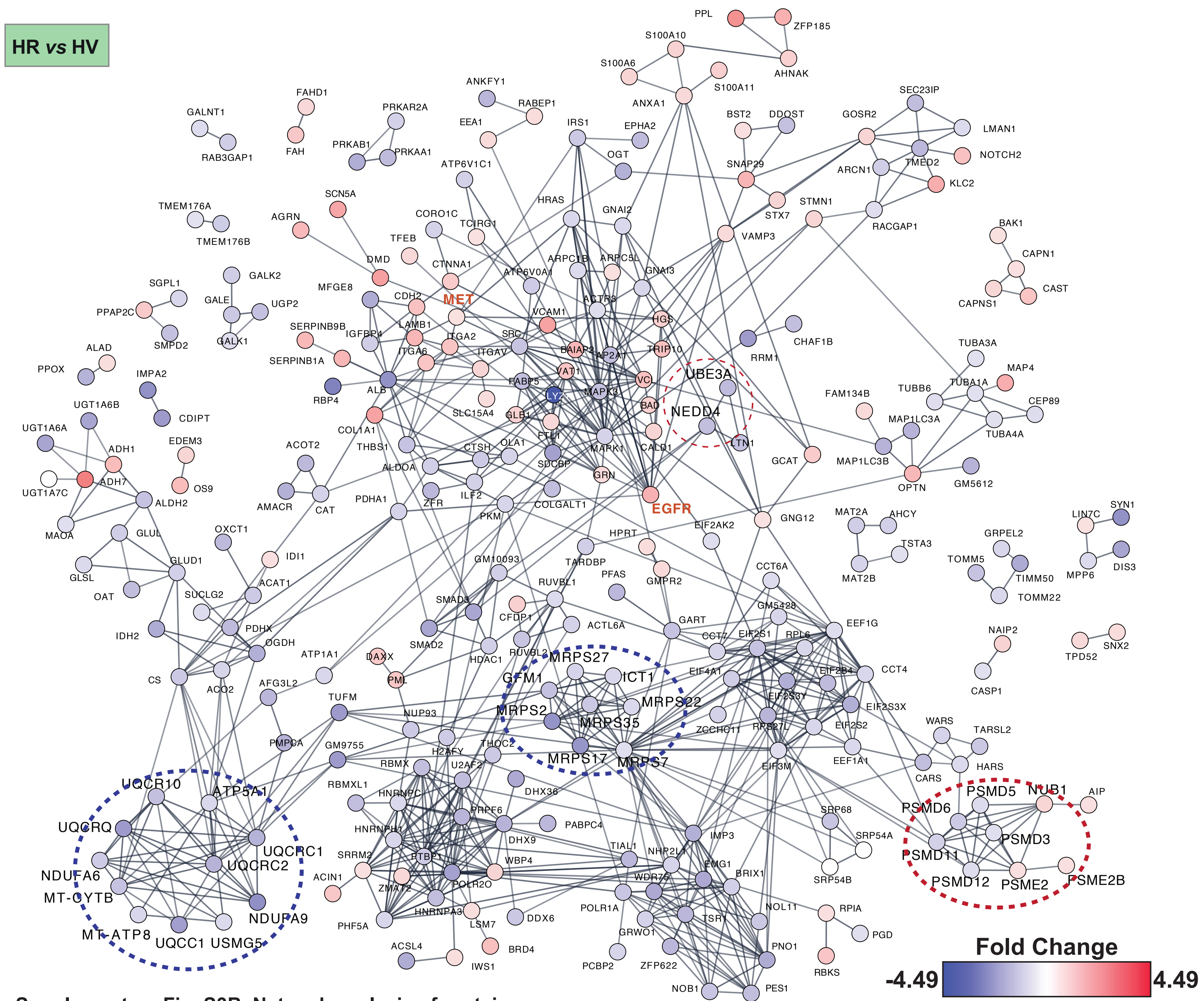


**Supplementary Fig. S3B. Network analysis of proteins modulated in HR cells.** Proteasome componenets are circled in red and mitochonrdial proteins in blue.

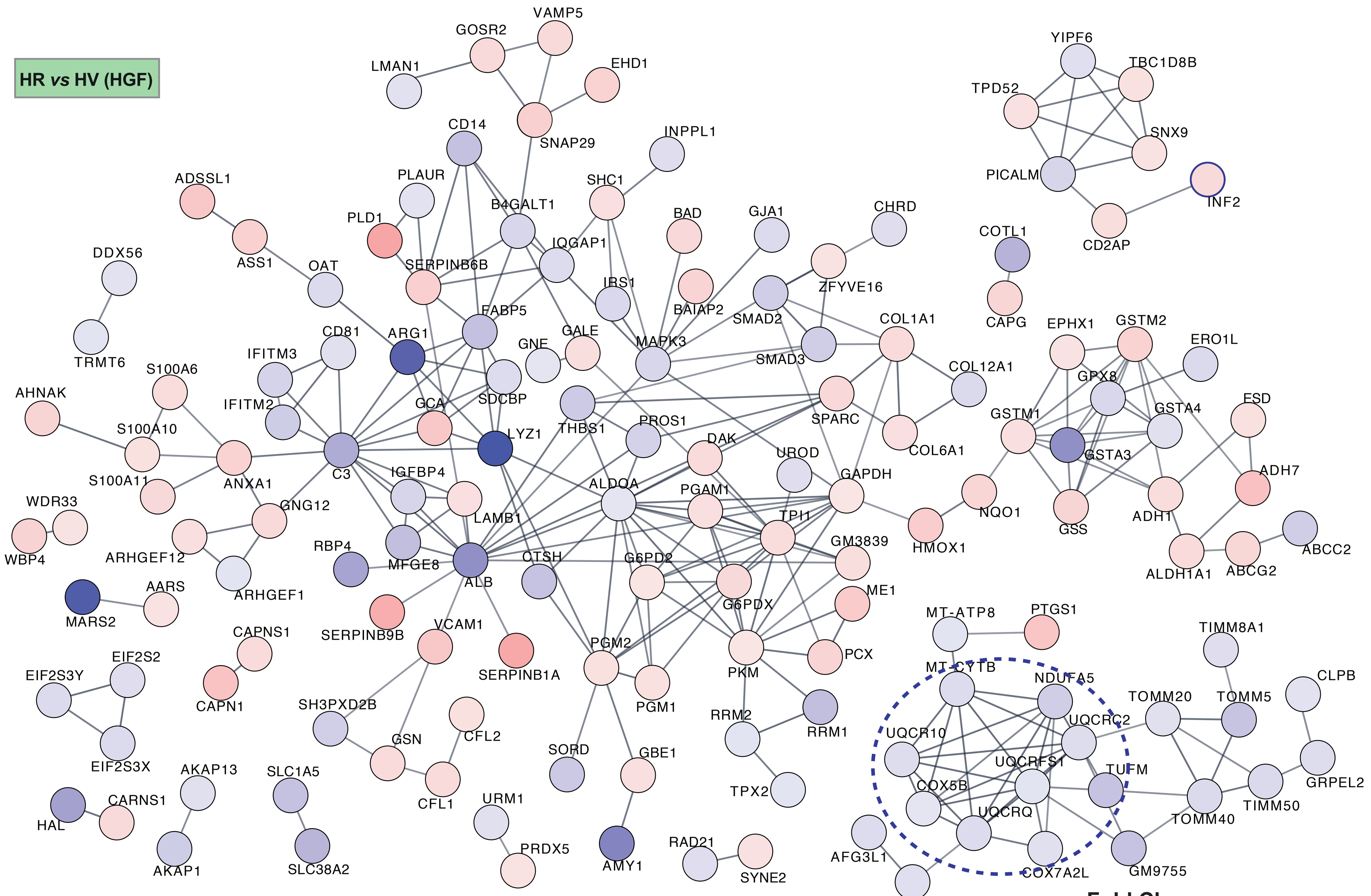


**Supplementary Fig. S3C. Network analysis of proteins modulated in HR cells following HGF stimulation.** Mitochonrdial proteins are circled in blue.
