## Supplementary Table S1 for "SILAC proteomics implicates the ubiquitin conjugating enzyme UBE2D in SOCS1-mediated downmodulation of the MET receptor in hepatocytes"

A) List of qRT-PCR primers used in this study.

| Gene Name | Accession Number | Forward Sequence (F) | Reverse Sequence (R) | Melting Temp $_o$C (F/R) | Amplicon Length (bp) |
|---|---|---|---|---|---|
| *Ube2d1* | NM_145420 | CCACCCAAATATAAACAGCAACG | TCTGGTACTAAGGGATCGTCTG | 62.3/62.4 | 144 |
| *Ube2d2* | NM_019912 | TCCTTTAGTGCCTGAGATTGC | AGTTCCCCTAGCTTTATTTGTAGAG | 62.2/62.1 | 140 |
| *Ube2d3* | NM_025356 | AGACTATGGCGCTGAAACG | GATATGGGCTGTCATTAGGTCC | 62.1/62.0 | 141 |
| *β-actin* | *NM_007393* | *ACCTTCTACAATGAGCTGCG* | *CTGGATGGCTACGTACATGG* | *62.1/61.8* | *147* |

B) List of western blot antibodies used in this study.

| Protein | Species | Source Company | Catalog Number | Dilution used |
|---|---|---|---|---|
| Phospo-STAT1 | Rabbit | Cell Signaling Technology | #9167 | 1:1000 |
| Total STAT1 | Rabbit | SantaCruz Biotech | sc-592 | 1:500 |
| Phospho-MET | Rabbit | Cell Signaling Technology | #3077 | 1:1000 |
| Total MET | Mouse | Santacruz Biotech | sc-8057 | 1:500 |
| c-CBL | Rabbit | Cell Signaling Technology | #2747 | 1:1000 |
| UBE2D | Mouse | Santacruz Biotech | sc-166278 | 1:500 |
| Actin | Rabbit | Cell Signaling Technology | #4970 | 1:1000 |
| HA tag | Mouse | SantaCruz Biotech | sc-7392 | 1:500 |
